# Atomic structure of the mitochondrial inner membrane AAA+ protease YME1 reveals the mechanism of substrate processing

**DOI:** 10.1101/189316

**Authors:** Cristina Puchades, Anthony J. Rampello, Mia Shin, Christopher J. Giuliano, R. Luke Wiseman, Steven E. Glynn, Gabriel C. Lander

## Abstract

We present the first atomic model of a substrate-bound inner mitochondrial membrane AAA+ quality control protease, YME1. Our ~3.4 Å cryo-EM structure reveals how the ATPases form a closed spiral staircase encircling an unfolded substrate, directing it toward the flat, symmetric protease ring. Importantly, the structure reveals how three coexisting nucleotide states allosterically induce distinct positioning of tyrosines in the central channel, resulting in substrate engagement and translocation to the negatively charged proteolytic chamber. This tight coordination by a network of conserved residues defines a sequential, around-the-ring ATP hydrolysis cycle that results in step-wise substrate translocation. Furthermore, we identify a hinge-like linker that accommodates the large-scale nucleotide-driven motions of the ATPase spiral independently of the contiguous planar proteolytic base. These results define the first molecular mechanism for a mitochondrial inner membrane AAA+ protease and reveal a translocation mechanism likely conserved for other AAA+ ATPases.

## Introduction

The regulation of mitochondrial protein quality control is essential for mitochondrial function and cellular survival. Imbalances in this regulation are closely associated with a variety of human diseases including many neurological and cardiovascular disorders (Nolden 2005, Tatsuta and Langer 2008, Quiros 2015). Since the majority of the mitochondrial proteome cannot be accessed by the cytosolic ubiquitin/proteasome pathways, mitochondrial protein quality control is primarily controlled by a network of proteases that degrade damaged or misfolded proteins (Leonhard 1996, Baker 2011), such as the AAA+ protease YME1L in the inner membrane (IM). YME1L is involved in nearly all aspects of mitochondrial biology, including regulation of the electron transport chain, protein import, lipid synthesis, and mitochondrial morphology (Leonhard 1999, Stiburek 2012, Potting 2013, Rainbolt 2013). Interestingly, stress-induced reductions in YME1L activity severely disrupt mitochondrial function and increase cellular stress-sensitivity both *in vitro* and *in vivo* (Baburamani 2015, Rainbolt 2015, Rainbolt 2016). Similarly, genetic ablation of *Yme1l1* is embryonic lethal and conditional deletion in adult cardiomyocytes causes heart failure and premature death in mice (Wai 2015). Furthermore, homozygous mutation of *Yme1l1* causes mitochondriopathy with optic nerve atrophy in humans (Hartmann 2016).

Despite its crucial function, a lack of structural information for YME1L or any other mitochondrial IM AAA+ protease significantly hinders our understanding of the mechanistic details that drive the proteolytic activity of these mitochondrial quality control machines. Unlike the proteolysis-associated AAA+ unfoldases found in the eukaryotic cytoplasm, which can interact with separate proteolytic complexes (i.e. the 20S core particle), mitochondrial IM AAA+ proteases contain both ATPase and protease domains on a single polypeptide, separated by a short linker region. In YME1L, the AAA+ ATPase domain and M41 peptidase domains reside in the mitochondrial intermembrane space (IMS), tethered to the IM by a single-pass membrane helix (Figure 1A,B). YME1L is evolutionarily related to bacterial FtsH and the catalytic domains exhibit a high degree of conservation across all eukaryotes (Koppen and Langer 2007, Quiros 2015). The catalytic domains of the yeast homolog, YME1, share 54% sequence identity with the human homologs, and expressing human *Yme1l1* in a *yme-1*-deficient yeast restores mitochondrial function, indicating comparable activities and substrate profiles (Shah 2000).

**Figure 1.**
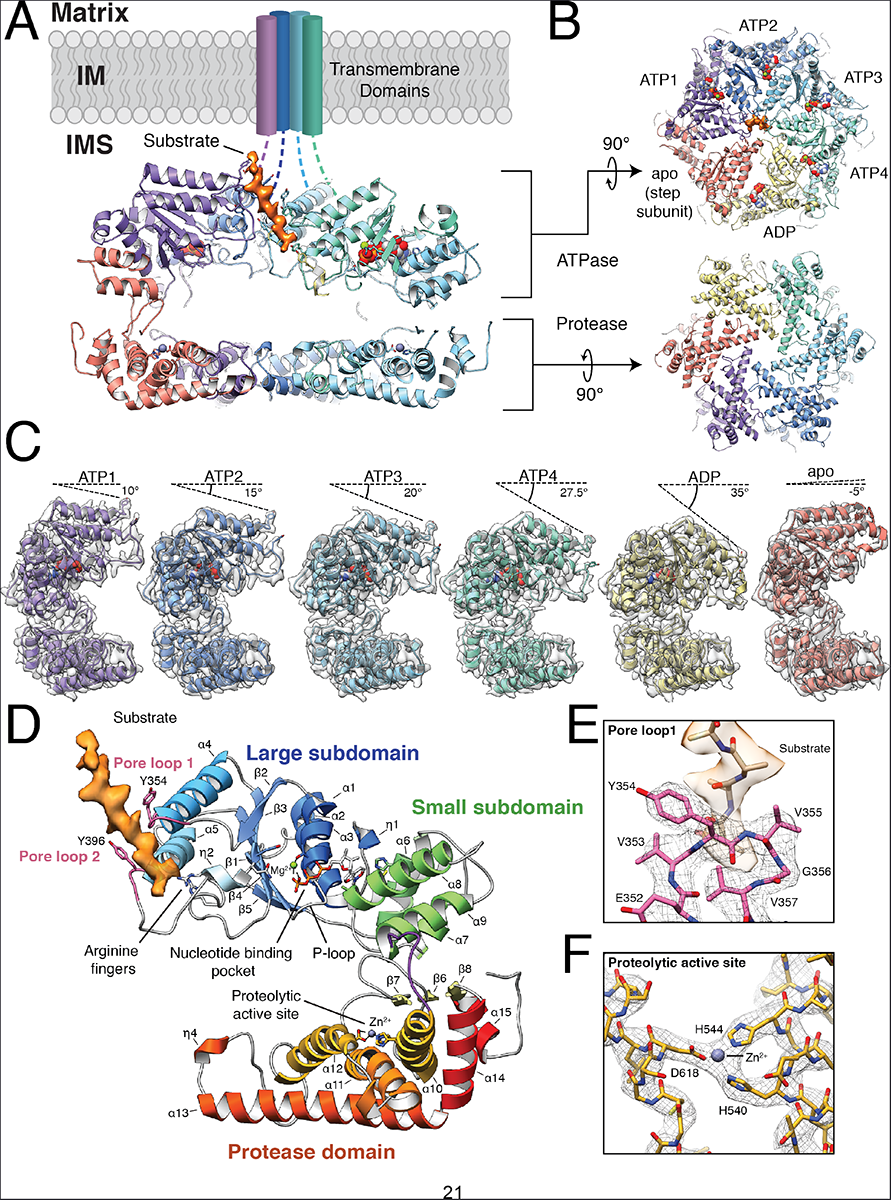
Architecture of the substrate-bound YME1 AAA+ protease. **A)** Cutaway view of the substrate-bound YME1 atomic model, with cryo-EM density for substrate colored orange. Each subunit of the homohexamer is assigned a unique color and the nucleic acids are depicted using a sphere representation in (**A-C**). **B)** Views of the ATPase and protease rings, shown orthogonally to the orientation shown in (A). **C)** Individual protomers shown side-by-side, aligned with the protease domain in the same orientation. The cryo-EM density is shown as a transparent gray isosurface, showing the quality of the reconstruction. The sequential movement of the ATPase domain relative to the protease domain is emphasized by a dashed line above each protomer, depicting the tilt of the pore loop helices relative to the horizontal protease ring. The pore loop tyrosines are visible within each of the pore loops. **D)** Topological organization of the YME1 protomer showing the large and small subdomains of the ATPase domain, and protease domain underneath. Notable and conserved components of the YME1 subunit are highlighted. The cryo-EM density of the substrate is shown in orange. **E)** Detailed view of the pore loop 1 interactions with substrate. The cryo-EM density of the substrate is shown as a transparent orange isosurface, along with a poly-alanine model of the substrate. The cryo-EM density of the pore loop is shown as a mesh, and the atomic model shows the interaction of Tyr354 and Val355 with substrate. **F)** Detailed view of the zinc-coordinated proteolytic active site.

A recently engineered soluble construct of yeast YME1, ^hex^YME1, comprising the AAA+ ATPase and protease domains assembled into active oligomers by fusion to a hexamerizing domain, exhibits high ATPase and proteolytic activity, as well as biologically relevant substrate specificity (Shi 2016, Rampello and Glynn 2017). Here, we employed a ^hex^YME1 variant containing an E381Q Walker B mutant (^hex^YME1^WB^) that limits ATP hydrolysis to enable structural determination of this ATP-dependent protease by cryo-electron microscopy (cryo-EM) (Figure 1, Supplementary Figures 1-2). We present the first atomic model of a mitochondrial IM AAA+ protease to reveal the allosteric mechanism linking ATP hydrolysis to substrate translocation in YME1.

### Quaternary organization of the substrate-engaged YME1 AAA+ protease

The ^hex^YME1^WB^ hexamer was solved in the presence of saturating amounts of ATP (1mM). The cryo-EM reconstruction was estimated to have an overall resolution of ~3.4 Å, with the best-resolved regions at ~3.2 Å, providing a near-atomic resolution view of the homohexameric nucleotide-bound ^hex^YME1^WB^ complex (Figure 1, Supplementary Figure 2, Supplementary Table 1). Our structure describes how the catalytic domains assemble into two stacked rings, with an asymmetric spiral staircase comprising the ATPase domains atop a planar, C6-symmetric protease ring (Figure 1A,B). Together, the ATPase and protease rings enclose a negatively charged proteolytic chamber encircled by the inter-domain linker (Figure 1A,B, Supplementary Figure 3A).

Previous crystal structures of FtsH in ADP-bound and nucleotide-free conformations contained ATPase domains organized into hexameric rings with 2-, 3-, and 6-fold symmetry (Suno 2006, Bieniossek 2009, Vostrukhina 2015). While the topological organization of the YME1 protomer closely resemble that of FtsH (Figure 1D), our structure reveals a distinctly different quaternary organization. Instead of a symmetric organization of the AAA+ domains, the YME1 AAA+ domains assemble into a spiral staircase with the ATPases progressively rotated and translated with respect to one another, similar to the organization observed for numerous other AAA+ unfoldases, including the functionally related 26S proteasome ATPase (Glynn 2009, Lander 2012, Matyskiela 2013, Monroe 2017, Ripstein 2017). In this arrangement, a “step” subunit (red in Figure 1) connects the lowest and highest positions of the staircase (yellow and purple in Figure 1, respectively). This step subunit exhibited high positional variability in our dataset, requiring focused 3D classification to enable confident atomic modeling of this protomer (Supplementary Figure 1D-G). However, this is the first description of this organization occurring for a AAA+ protease where the ATPase and protease domains are contained within the same polypeptide chain. Moreover, the resolution of our structure, which is close to ~3 Å for much of the reconstruction (Supplementary Figure 2), enables us to explore the atomic features of the staircase architecture in greater detail than these previous studies (Figure 1E).

The central pore of the ATPase spiral staircase has a diameter of ~1.4 nm, which is sufficient to accommodate an unfolded peptide (Figure 1B). Surprisingly, we discovered an additional density within the pore, oriented at a 28º angle relative to the hexameric y-axis, into which an unfolded 10 amino acid peptide could be modeled (Figure 1A,B,D, movie 1). We conclude that this density represents a substrate peptide that is trapped in the process of translocation through the YME1 pore. Cryo-EM reconstructions of other ATPases in similar asymmetric spiraling configurations also revealed substrate density in the central pores (Gates 2017, Monroe 2017, Ripstein 2017), suggesting that this conformation may be induced by substrate binding. The presence of this substrate and the resolution of our structure provide a unique opportunity to define how YME1 handles substrates during the translocation process.

### YME1 ATPase domains engage substrates through a double spiral staircase of tyrosines

A conserved aromatic-hydrophobic motif (typically Tyr/Phe-Val) in the central pore loop 1 is found across AAA+ ATPases and numerous studies show direct involvement of the motif in AAA+-dependent unfolding and translocation (Schlieker 2004, Burton 2005, Hinnerwisch 2005, Park 2005, Okuno 2006, Martin 2008). In YME1, mutation of the conserved aromatic residue (Y354) impairs substrate degradation, confirming the importance of this residue (Graef and Langer 2006). We visualize direct interactions between Y354 and the translocating substrate in our YME1 complex, in which the Y354 residues adopt a spiral staircase organization that mirrors the global architecture of the AAA+ domains (Figure 1E, 2A). Interestingly, the Y354 residues show three distinct modes of interaction with the translocating substrate, relative to their position in the staircase architecture. The central four Y354 residues are similarly intercalated into the substrate, with each Y354 positioned between substrate side chains to engage the substrate backbone in a zipper-like configuration (Figure 2A, Supplementary Figure 4), a non-specific mode of interaction that is compatible with sequence-independent substrate translocation essential for the function of YME1. However, the lowest Y354 (yellow in Figure 2A) shows a more modest interaction with substrate, and Y354 of the step subunit, whose pore-loop 1 adopts the top-most position within the Tyr staircase (red in Figure 2A), is completely disengaged from the substrate. This indicates that the positioning of the specific ATPases within the spiral staircase dictates the engagement of Y354 with substrate. Additionally, our structure shows that the adjacent valine of the conserved aromatic-hydrophobic motif of pore-loop 1 (V355) is also directed towards the substrate side chains (Figure 1E), in agreement with previous biochemical data implicating this residue in dictating substrate specificity for FtsH (Yamada-Inagawa 2003, Okuno 2006).

**Figure 2.**
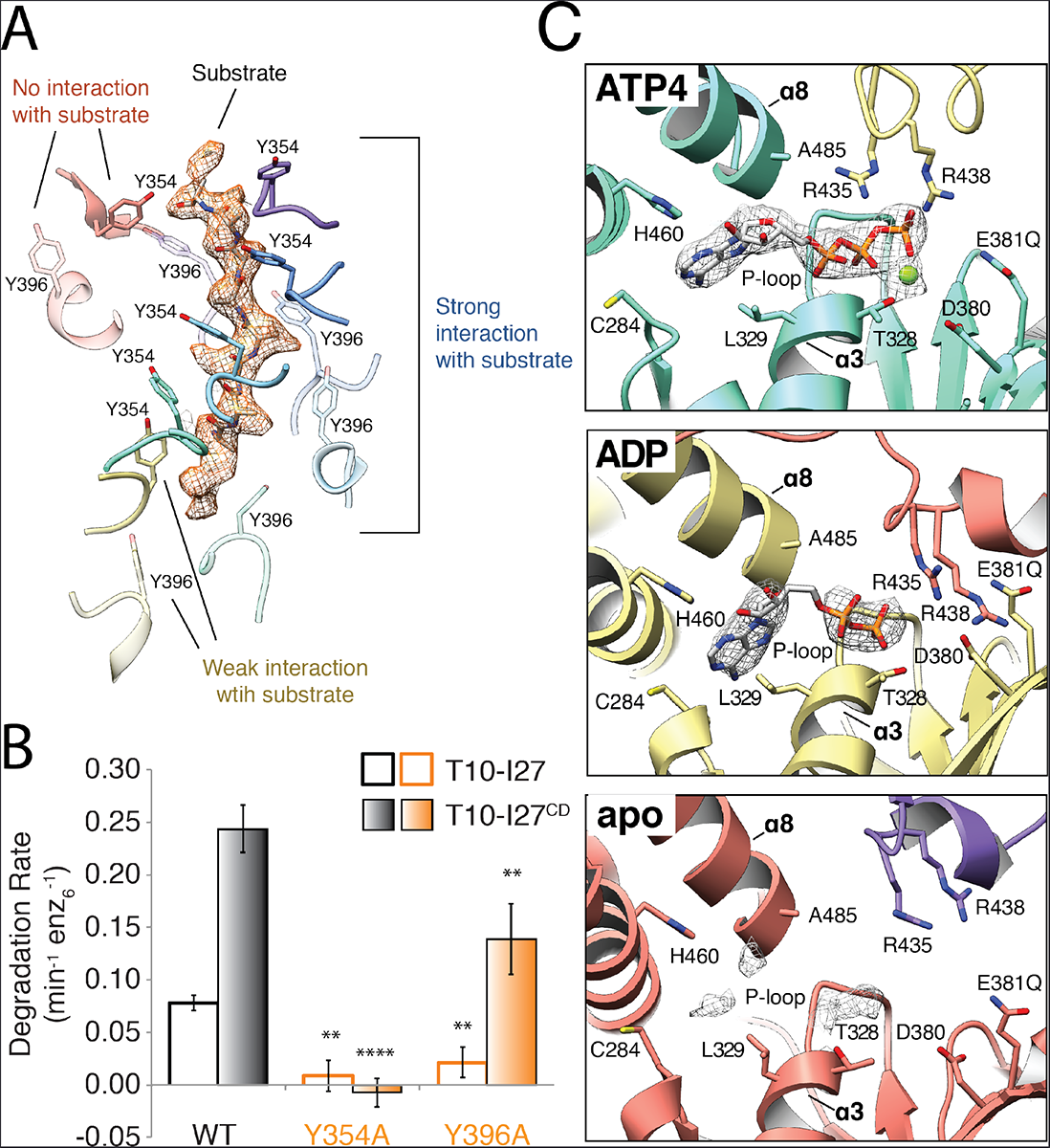
Nucleotide state correlates with pore loop conformation and substrate interaction. **A)** Three categories of interactions are observed between the pore loop tyrosines and substrate. A poly-alanine model of the substrate is shown within the cryo-EM density of the substrate, shown as orange mesh. Tyr354 and Tyr396 from pore loops 1 and 2 are rendered as solid and semi-transparent, respectively. The Tyr residues from the ATP(1-4) subunits (purple, blue, cyan, and turquoise) show close, stable, intercalating interactions with substrate, while Tyr residues from the ADP subunit (yellow) are positioned further from the substrate, and Tyr residues from the apo subunit (red) show no interaction with substrate. **B)** Effect of mutations in the conserved pore loops on substrate degradation. Initial degradation rates are shown for folded (T10-I27) and unfolded (T10-I27^CD^) substrates by wild-type ^hex^YME1 and variants bearing either Y354A or Y396A substitutions. Mutation of the pore loop 1 Tyrosine abolishes degradation for both substrates and mutation of the pore loop 2 Tyrosine significantly diminishes the degradation rate for both substrates, consistent with an important role for both of these residues in substrate handling. Values are means of independent replicates (n≥3) ± s.d. \*\**P*≤0.01, \*\*\*\**P<*0.0001 as calculated using the Student’s two-tailed t-test and shown in comparison to the degradation of the identical substrate by wild-type ^hex^YME1. **C)** Organization of the nucleotide binding pockets in the ATP(1-4), ADP, and apo states. Cryo-EM density corresponding to the nucleotide (or lack thereof) is shown as a gray mesh at a σ level of 4.0 for all states. Absence of gamma phosphate and magnesium ion is evident in the ADP-like state, whereas only a low level of nucleotide density is still present in the apo-conformation (bottom panel). In the ATP(1-4) and ADP conformations, the adenine base is located within 4 Å of L329 and C284 of the large domain and H460 of the small domain, whereas the ribose directly interacts with the backbone of A485 from helix **a**8 of the small domain. The arginine fingers (R435 and R438) are closely coordinated with the phosphates in the ATP(1-4) subunits, but further away in the ADP and apo subunits.

Unexpectedly, we also observed that another tyrosine residue, Y396 within the pore loop 2 of the YME1 AAA+ domains, forms a second spiral arrangement surrounding the substrate at a position below the pore loop 1 staircase (Figure 2A). The interactions between Y396 and the translocating substrate mirrors those observed for the pore-loop 1 Y354, wherein the Y396 residues within the central four ATPase domains in the spiral staircase appear to interact with substrate, while the Y396 of the lowest subunit shows a weaker interaction with the substrate and the Y396 of the step subunit completely is disengaged. While the presence of a tyrosine residue in this position is conserved across YME1 homologs, including mammalian YME1L, this residue is not found in more distantly related mitochondrial enzymes such as AFG3L2 or in cytosolic AAA+ proteases such as the 26S proteasome.

To confirm that these observed pore-loop interactions play a role in substrate translocation, we examined the substrate degradation activity of ^hex^YME1 containing point mutations in the pore-loop 1 tyrosine (Y354A) or the pore-loop 2 tyrosine (Y396A). Introduction of these mutations into either pore loop largely abolish degradation of the I27 domain of human titin bearing an N-terminal degron derived from the mitochondrial Tim10 protein (T10-I27) (Figure 2B) (Rampello and Glynn 2017). The introduction of mutations in the I27 domain to destabilize the folded structure (T10-I27^CD^) produced a large increase in degradation rate by wild-type ^hex^YME1, likely resulting from the removal of the unfolding step of degradation (Kenniston 2003). This unfolded protein was not degraded by ^hex^YME1 bearing the Y354A mutation, and the degradation rate of Y396A was significantly reduced compared to wild-type. Together, these results demonstrate that both pore loop tyrosines are important for substrate degradation. Interestingly, the Y354A mutant, but not Y396A, also impaired ATP hydrolysis, suggesting allosteric coordination linking substrate interactions at the pore-loop with the nucleotide binding pocket (Supplementary Figure 1B). This phenomenon was previously shown biochemically for FtsH and other AAA+ ATPases (Yamada-Inagawa 2003, Zhang and Wigley 2008, Chang 2017), but the mechanism of this allostery has remained elusive, prompting us to explore the relationship between nucleotide state and the observed pore-loop conformations.

### YME1 contains three co-existing nucleotide states

The resolution of our reconstruction enabled us to unambiguously define the nucleotide state for each subunit within the YME1 homohexamer. We observe three distinct nucleotide states co-existing within the structure. Four subunits in the YME1 AAA+ spiral staircase (ATP1, ATP2, ATP3, and ATP4 shown in Figure 1B) contain a well-resolved ATP in the nucleotide binding pocket, clearly defined by the presence of the three phosphate residues (Figure 2C and Supplementary Figure 5A,B, movie 1). The lowest subunit of the spiral staircase was found to have an ADP-like density in the nucleotide binding pocket (ADP in Figure 1B, Figure 2C, and Supplementary Figure 5A,B, movie 1), whereas the binding pocket of the step subunit contains weak nucleotide density, which we refer to as “apo-like” (Figure 2C, Supplementary Figure 5A,B, movie 1). The presence of ADP-like density in the lowest subunit can be explained by the presence of small amounts of contaminating ADP or by low residual ATP hydrolysis in the Walker-B mutant (Hersch 2005, Beckwith 2013).

Importantly, these three distinct nucleotide states correlate directly with the three modes of interaction we observe between the pore-loop tyrosines and the translocating substrates. The pore loop tyrosines of the central four subunits from the staircase (ATP1-4), which all directly engage the substrate in an identical fashion, belong to subunits bound to ATP (Figure 1A,B, Figure 2A). The tyrosine residues at the bottom of the staircase, which show a weaker interaction with substrate, belong to the ADP-bound subunit, while the pore-loop tyrosines that are completely disengaged from the substrate at the top of the staircase, belong to the apo-like step subunit. This direct correlation between nucleotide states and substrate engagement, combined with our biochemical results, indicate that nucleotide hydrolysis and release provides a mechanism to allosterically regulate the pore loop conformations and direct substrate translocation.

### The spiral staircase organization links nucleotide state to pore loop conformation

Nucleotides bind to AAA+ ATPases within a pocket formed at the interface of the large and small subdomains of one ATPase and the large subdomain of the neighboring ATPase (Figure 2C). YME1 belongs to the P-loop subclass of ATPases, which are characterized by a conserved Walker A motif (GXXGXGK[S/T]) in the large subdomain that directly interacts with nucleotide (Hanson and Whiteheart 2005). Accordingly, backbone nitrogens of the P-loop residues G324 and G326 directly interact with the nucleotide phosphates in our YME1 structure (Supplementary Figure 5B). Notably, all residues that were previously found to be crucial for nucleotide binding in YME1 (Gerdes 2012) are involved in nucleotide interaction in our structure (Figure 2C).

The machinery of ATP hydrolysis is highly conserved across AAA+ ATPases, simultaneously involving side chains from both the large and small ATPase domains, as well as trans-acting residues of the neighboring subunit. Within the YME1 nucleotide-binding pocket, a threonine (T328 in α 3 of YME1) coordinates a magnesium ion with the ATP phosphates (Figure 2C, top panel, and Figure 3A). In addition, two consecutive acidic residues (D380 and E381 in YME1 β1) coordinate and activate a catalytic water molecule. The large ATPase subdomain of the neighboring protomer contributes to the coordination of the ATP gamma phosphate through two highly conserved “arginine fingers” (R435 and R438 in YME1) *(Hanson and Whiteheart 2005)*. Furthermore, in the presence of ATP, we observe a loop (D409-G410-F411) at the C-terminal end of α5 of this adjacent subdomain that bridges across the nucleotide-binding pocket, contacting the ATP-bound subunit. Notably, this loop is strictly conserved across homologs and paralogs of YME1, as well as in the ATPases of the 26S proteasome (Supplementary Figure 6A). In fact, this region has been previously identified as the inter-subunit signaling (ISS) motif, as mutagenesis of these residues revealed a critical role in communicating nucleotide state between protomers (Augustin 2009). Importantly, in this extended conformation of the ISS motif, D409 is in close proximity to the Arginine fingers and F411 packs against phenylalanine residues in β2 and β3 of the ATP-bound subunit (Figure 3A, purple in left panel, movie 2). These aromatic interactions position the pore loop 1 and 2 tyrosines protruding from α4 and α5 of the ATP-bound subunit in an orientation that is compatible with substrate engagement (Figure 3A, pink in left panel, Figure 3B, green, and Supplementary Figure 7).

**Figure 3.**
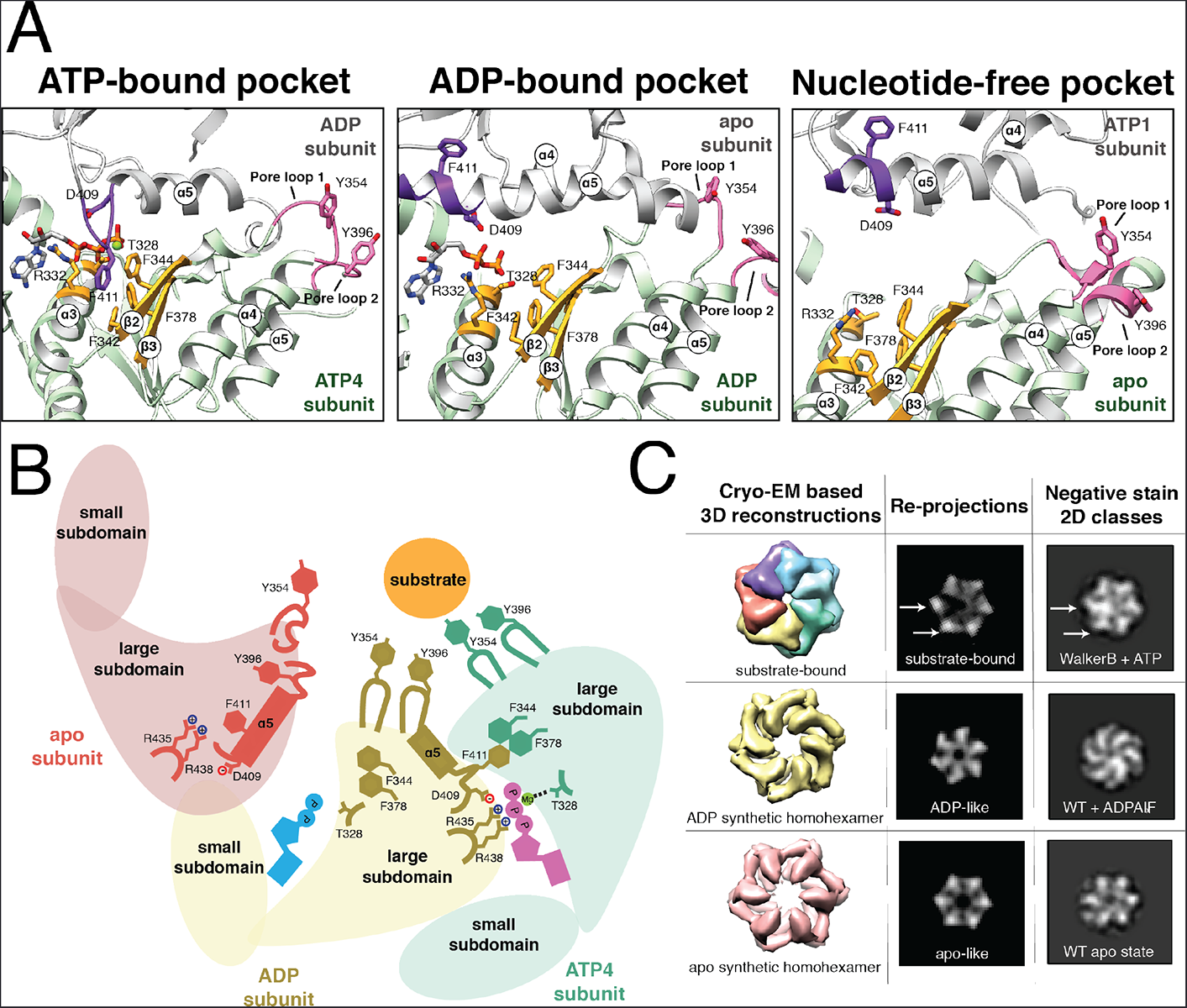
Nucleotide state causes major conformational rearrangements of the ATPase domain. **A)** The three nucleotide state conformations were aligned to the central phenylalanine-containing β-sheet (colored gold) to show the major rearrangements that allosterically affect pore loop conformation. In the ATP-bound pocket, the ISS loop of the adjacent subunit (colored purple) extends towards the Phe-containing β-sheet. In this conformation, the pore loops (colored pink) are extended toward the central pore and interact strongly with substrate. In the ADP-bound pocket, loss of the gamma phosphate results in a retraction of the ISS loop from the Phe-containing β-sheet, and the ISS loop becomes part of the α5 helix of the adjacent subunit. Within the ADP-bound subunit, the α3 helix moves closer to the Phe-containing β -sheet, and the pore-loop tyrosines more weakly interact with substrate. In the nucleotide-free state, the pore loop tyrosines are completely dissociated from the substrate, and are incorporated into helices 4 and 5. **B)** Cartoon representation of the nucleotide-dependent conformational changes affecting the entire ATPase domain, depicting the motions described in (A). When ATP is bound within the nucleotide pocket, the pore loop tyrosines interact strongly with substrate. ATP hydrolysis results in a weakening of the substrate-tyrosine interactions, and loss of the nucleotide results in a complete release of substrate. **C)** Nucleotide-state is shown to cause major domain rotations, as 3D structures of the substrate-bound and computationally symmetrized ADP and apo homohexamers generate distinctly different 2D projections that show structural characteristics that are similar to those in reference-free 2D classes obtained by negative stain EM in the presence of ATP, ADPALF_X_ or in the absence of nucleotide. Wild type ^hex^YME1 is abbreviated as WT.

Loss of the ATP gamma phosphate disrupts the inter-protomer coordination between the ISS motif in the apo step subunit and the nucleotide binding pocket within the ADP-like subunit (Figure 3A,B and Supplementary Figure 7). This results in a two-fold increase in the distance between the step subunit’s arginine fingers and the ADP-like nucleotide-binding site. Interestingly, this reduced coordination between subunits causes the ISS motif of the apo step subunit to retract from the neighboring nucleotide binding pocket of the ADP-bound subunit and become incorporated into the C-terminal end of a now extended α5 (Figure 3A, purple in middle panel, movie 2). This helical refolding of the ISS loop disrupts the aromatic interactions between F411 of the apo step subunit and the phenylalanine residues in the β2 and β3 strands of the ADP-like subunit. In the absence of the trans-acting F411 from the apo step subunit, the α3 helix of the ADP-like subunit is drawn closer to the central β-sheet, shifting T328 away from the magnesium coordination site. Together, these rearrangements reconfigure the pore-loop tyrosines in the ADP subunit to orientations positioned away from the substrate (Figure 3A, pink in middle panel and Figure 3B, brown, and Supplementary Figure 7).

As described above, the apo step subunit contains a ISS motif folded into the C-terminus α5 helix, precluding interaction between F411 and the ADP-bound subunit (Figure 3A, purple in middle panel, Supplementary Figure 7, movie 2). In addition, the lack of nucleotide in the step subunit itself prevents inter-protomer stabilization by a trans-acting F411 from the ATP1 subunit (Figure 3A, purple in right panel, Supplementary Figure 7). As a result, the step subunit is disconnected from both neighboring protomers, explaining the high positional flexibility for this subunit observed in our cryo-EM reconstruction. Interestingly, this lack of inter-protomer coordination results in an N-terminal extension of the step subunit α5 helix, altering the conformation of pore-loop 2 and resulting in its incorporation into the α5 helix (Fig 3A, pink in right panel, Figure 3B). The extension of the α5 helix at both the N- and C-terminal ends results in a sequestration of the pore-loop 2 Y396 from substrate interaction, and simultaneously influences the conformation of the adjacent α4 helix to reposition the pore-loop 1 Y354 away from the substrate (Supplementary Figure 7).

Taken together, our results indicate that nucleotide binding and hydrolysis regulate the interactions between the pore-loop tyrosine double staircase and the substrate (Figure 3B and Supplementary Figure 7). In the ATP-bound subunits, the nucleotide binding pocket is positioned to allow hydrolysis, and while the precise trigger for hydrolysis is not clear from the structure, it is likely that subtle changes in the position of the arginine fingers from the adjacent ADP-like subunit are required to trigger hydrolysis in the ATP4 binding site. The structure clearly shows that the loss of the gamma phosphate induces remodeling of the nucleotide binding pocket, which reorganizes the pore-loop tyrosine residues of both the ADP-like subunit and the neighboring step subunit, dissociating these subunits from the substrate. Thus, the three nucleotide states coexisting in our substrate-bound structure strongly suggest that nucleotide state allosterically regulates pore loop interactions with substrates through conformational remodeling of the AAA+ ATPase domain.

### Nucleotide state coordinates major structural rearrangements within the ATPase domains

To better define the inter-domain reorganizations induced by nucleotide state, we calculated difference distance matrix plots that depict changes in distances between Cα atoms between one subunit and its clockwise neighbor (Supplementary Figure 8A). These matrices clearly show that the small and large subdomains undergo substantial rigid-body movements independently of each other in a nucleotide-dependent manner. Interestingly, the subdomains are closest to one another in the ADP-like subunit, whereas the distance is maximal in the step subunit (Supplementary Figure 8A).

To investigate the role that nucleotide state plays in directing this subdomain positioning, we structurally characterized ^hex^YME1^WT^ in the absence of nucleotide, as well as in the presence of ADPAlF_X_, an ATP analog known to induce ATP-hydrolysis-intermediate states in other AAA+ ATPases (DeLaBarre and Brunger 2005, Han 2015). Notably, in both the absence of ATP and presence of ADPAlF_X_, the conformational heterogeneity of the hexamer rose to a level that impeded 3D reconstructions. However, 2D analysis using negative stain EM was sufficient to inform on the overall domain organization of these hexameric states. Reference-free 2D analyses revealed three remarkably different conformations in the absence of nucleotide or in presence of ATP and ADPAlF_X_, confirming that a reorganization of the protomer is associated with specific nucleotide states (Figure 3C, right column).

To assess whether the distinct tertiary organizations represented by these 2D analyses are consistent with the nucleotide-induced rearrangements we observe in our cryo-EM reconstruction, we used our atomic model to generate 3D densities representing each of the nucleotide states we resolved by 2D analyses. These densities were then filtered to a resolution comparable to negative stain. 2D projections of the filtered substrate-bound ^hex^YME1^WB^ were consistent with the negative stain 2D averages of this construct in the presence of ATP, serving as a positive control for these analyses (Figure 3C, top row). Importantly, the negative stain data showed that two of the protomers were in notably different conformations from the other 4 subunits in the complex, consistent with the presence of three distinct nucleotide states within our 3D structure (Figure 3C, top row arrows). In contrast, our 2D analyses showed that all subunits in the apo and ADPAlF_x_ samples adopted similar conformations within the hexamer, indicating that in each case the ATPases were in a homogeneous nucleotide state. The projections of the homohexamers generated from six copies of the ADP-like and apo-like subunits revealed a structural organization that accurately reflects the ADPAlF_x_–bound and apo negative-stain 2D classes, respectively. Serving as further confirmation for these nucleotide-induced reorganizations, symmetric crystallographic structures of FtsH in the apo and ADP-bound state also resemble our apo and ADP-like synthetic homohexamers (Bieniossek 2009, Vostrukhina 2015). Together, these data confirm the coexistence of three distinct nucleotide states in our substrate-bound structure, and show that nucleotide-state is sufficient to induce major rotations of the ATPase subdomains in YME1. Moreover, our results strongly suggest that these nucleotide-dependent motions are required for formation of the asymmetric spiral staircase configuration, as the subdomain movements induced by the ADP-like and apo-like nucleotide state place these subunits in the lower-most and step positions, respectively.

### A hinge-like glycine linker accommodates ATPase rearrangements above a flat protease ring

Notably, the considerable movements of the ATPase subdomains do not involve a reorganization of the protease domains, which maintain a C6-symmetric planar organization (Figure 1A-C). As a result, the distance between the ATPase and protease domains is gradually reduced as the protomers progress around the ring (Figure 1C, movie 1), which is further illustrated by distance matrices of the entire subunit (Supplementary Figure 8B). We hypothesized that the inter-domain linker connecting the protease and ATPase domains of each protomer plays a crucial role in accommodating the ATPase domain motions while adhered to a planar protease ring. We noted that a residue (G521) within the linker region enables the small ATPase domain to undergo a hinge-like motion relative to the protease domain (Figure 4A). This glycine is strictly conserved from bacteria to human, and allows for the range of phi and psi angles for the different nucleotide states observed in our YME1 structure (Figure 4B). Furthermore, previous biochemical and structural studies of FtsH highlighted the importance of this glycine for protease activity (Bieniossek 2009). To determine the structural and functional implications of this inter-domain hinge for YME1 function, we mutated this glycine to a leucine to limit the adoptable backbone dihedral angles (G521L). Negative stain EM analysis revealed that, regardless of nucleotide state, the mutant construct always adopted an ADP-like conformation (c.f., Figure 3C and Figure 4C). These results indicate that the large nucleotide-induced motions of the ATPase domains are enabled by the wide range of backbone conformations conferred by the dihedral properties of the inter-domain glycine. Accordingly, this point mutation completely abolished YME1’s ability to degrade the unfolded substrate T10-I27^CD^ despite hydrolyzing ATP at ~70% of the wild type rate (Figure 4D, Supplementary Figure 1B). This indicates that ATP hydrolysis is required, but not necessarily sufficient, for substrate processing, as the ATPase domain rotations enabled by the glycine linker are required for substrate translocation and ultimately proteolytic degradation.

**Figure 4.**
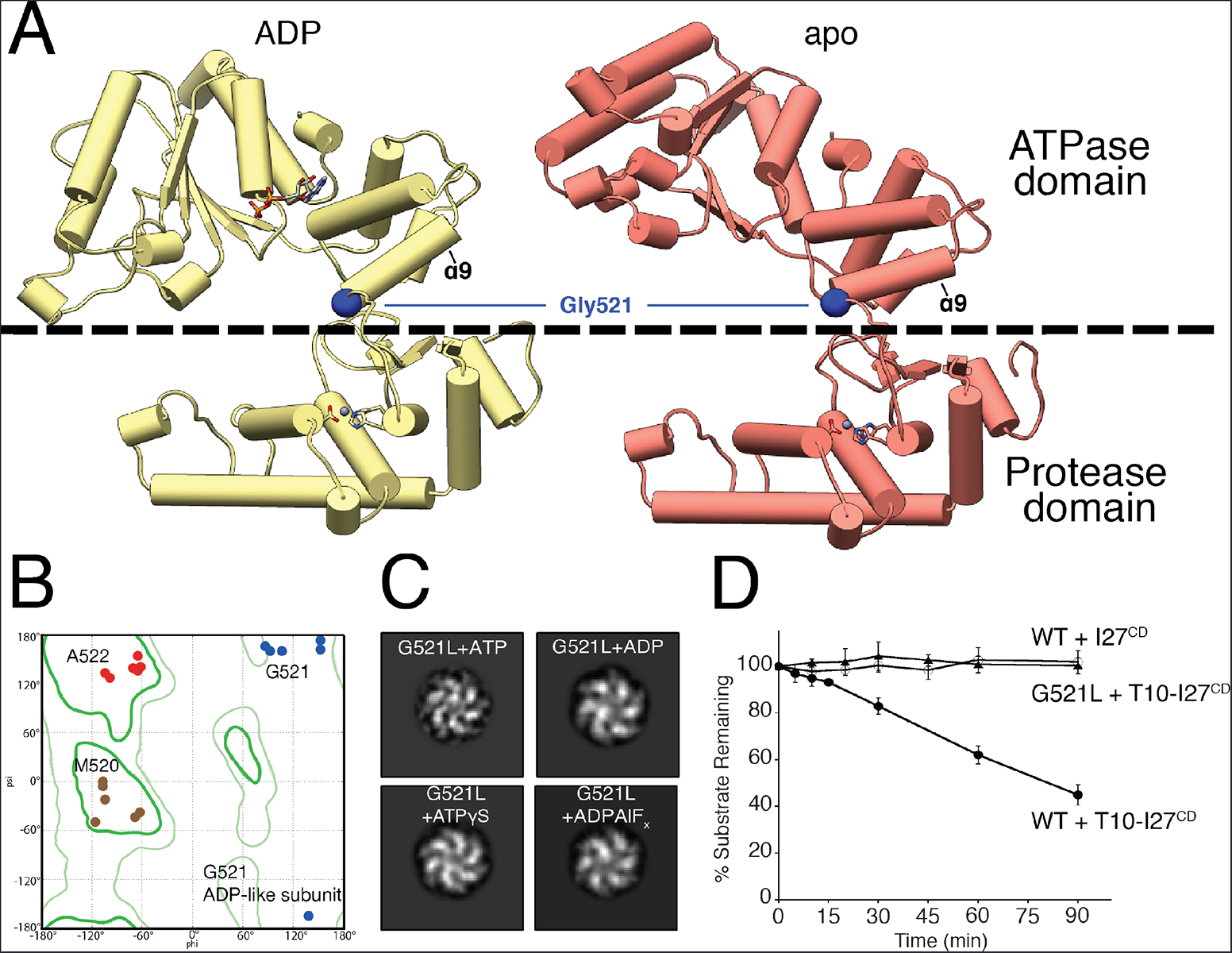
A hinge-like glycine linker accommodates nucleotide-state induced domain rotations. **A)** The ADP and apo subunits are shown (yellow and red, respectively), aligned relative to their protease domains. The dashed line delineates the ATPase domain (above) from the protease domain (below). The ATPase domain undergoes a large movement as it progresses from the ADP to the nucleotide-free state, which stems from a pivoting of the α9 helix around the inter-domain linker. Gly521 within this linker is shown to play a key role in enabling this large-scale motion. **B)** Ramachandran plot displaying phi and psi angles of inter-domain residues M520 (brown), G521 (blue) and A522 (red) in each subunit shows major torsion changes of G521 in the ADP-like state. The location of the Gly521 phi and psi angles underscore the necessity of a torsionally flexible Gly in this position. **C)** 2D image analyses of a negatively stained ^hex^YME1 construct containing a G521L mutation show that addition of ATP, ADP, ADPALF_X_, and ATPγS do not significantly influence the quaternary organization of the complex. Since all the resulting class averages appear to have a similar organization as the ADP-like conformation shown in Figure 3C, we suspect that the subunits of this mutant construct are trapped in the ADP conformation. **D)** Plot showing the effect of the G521L mutation on the degradation of an unfolded substrate. Rapid loss of T10-I27^CD^ is seen over time in the presence of wild-type ^hex^YME1 whereas no loss of T10-I27^CD^ is observed in the presence of the G521L point mutant or for unfolded I27^CD^ lacking the T10 degron incubated with wild-type ^hex^YME1. Values are means of independent replicates (n≥3) ± s.d.

### Positioning of the unfolded peptide in the protease domain can enable processive degradation of substrate

Our results show that the ATPase domain movements facilitate the translocation of an unfolded substrate peptide through the hydrophobic central pore into the hydrophilic chamber, where the protease domains comprise a very negatively charged ring with all 6 cleavage sites facing the interior of the chamber (Supplementary Figure 3A). The proteolytic active site of YME1 is well resolved in the reconstruction, showing how a zinc ion is coordinated by two histidines (H540 and H544) and an aspartate (D618), (Figure 1F) in an organization that is similar to that observed for other M41 proteases, such as the homologous FtsH (Suno 2006, Bieniossek 2009, Vostrukhina 2015). The active site is found at the periphery of the protease ring in a small pocket that constitutes the only hydrophobic patch accessible from the interior of the chamber (Supplementary Figure 3B). An anti-parallel β-strand is located directly adjacent to the proteolytic pocket, and we show that a ten amino acid poly-alanine unfolded substrate peptide can interact with this β-strand at the active site, precisely positioning it for proteolytic cleavage (Supplementary Figure 3C,D). The formation of interactions with the substrate polypeptide backbone atoms could enable sequence-independent cleavage, compatible with degradation of the wide variety of YME1 substrates identified *in vivo* (Graef and Langer 2006, Graef 2007, Potting 2010, Wang 2013).

### Model for the mechanism of action of YME1 and implications for other AAA+ unfoldases

We can use our data to define an ATP-dependent mechanism of substrate translocation by YME1. We show that nucleotide-state determines inter-protomer coordination, ATPase subdomain motions, and pore-loop conformation (Figure 5, A and B). Firstly, ATP hydrolysis in the lower-most ATP-bound subunit abolishes the interaction of the gamma phosphate with the trans-acting arginine finger residues from the clockwise adjacent subunit, thus releasing the bridging F411 and breaking the subunit-subunit interaction. This drives a major domain rotation that repositions this subunit in the lower-most register of the spiral staircase and weakens the interaction of its pore loop tyrosines with the substrate. These motions then trigger ATP hydrolysis in the counterclockwise adjacent subunit, likely through subtle repositioning of the arginine fingers that are coordinating the neighboring ATP. Evidence of this coordination of nucleotide hydrolysis is seen in a study of YME1 paralogs, Yta10 and Yta12, which showed that ATP binding within a given subunit inhibits ATP hydrolysis in the counterclockwise adjacent subunit (Augustin 2009). As the counterclockwise subunit subsequently undergoes hydrolysis, its contacts with the triggering ADP-like subunit are lost, leaving the ADP-like subunit now untethered from both neighboring subunits. As a result, the ADP-like subunit becomes displaced from the hexamer and releases ADP, thereby transitioning to an apo-like step subunit state and completely breaking interaction of the pore loops with the substrate. ATP binding by the apo step subunit reestablishes the interactions between the gamma phosphate and the trans-acting elements of its clockwise adjacent ATP-bound subunit to restore the spiral staircase arrangement with the previous apo subunit now occupying the higher-most ATP-bound position. Iteration of this sequence of events leads to a sequential ATP-hydrolysis cycle that proceeds around the ring in a counterclockwise manner (Figure 5A). This sequential model is in agreement with mechanisms proposed for other AAA+ ATPases (Enemark and Joshua-Tor 2006, Augustin 2009, Hoskins 2009, Gates 2017, Monroe 2017, Ripstein 2017).

**Figure 5.**
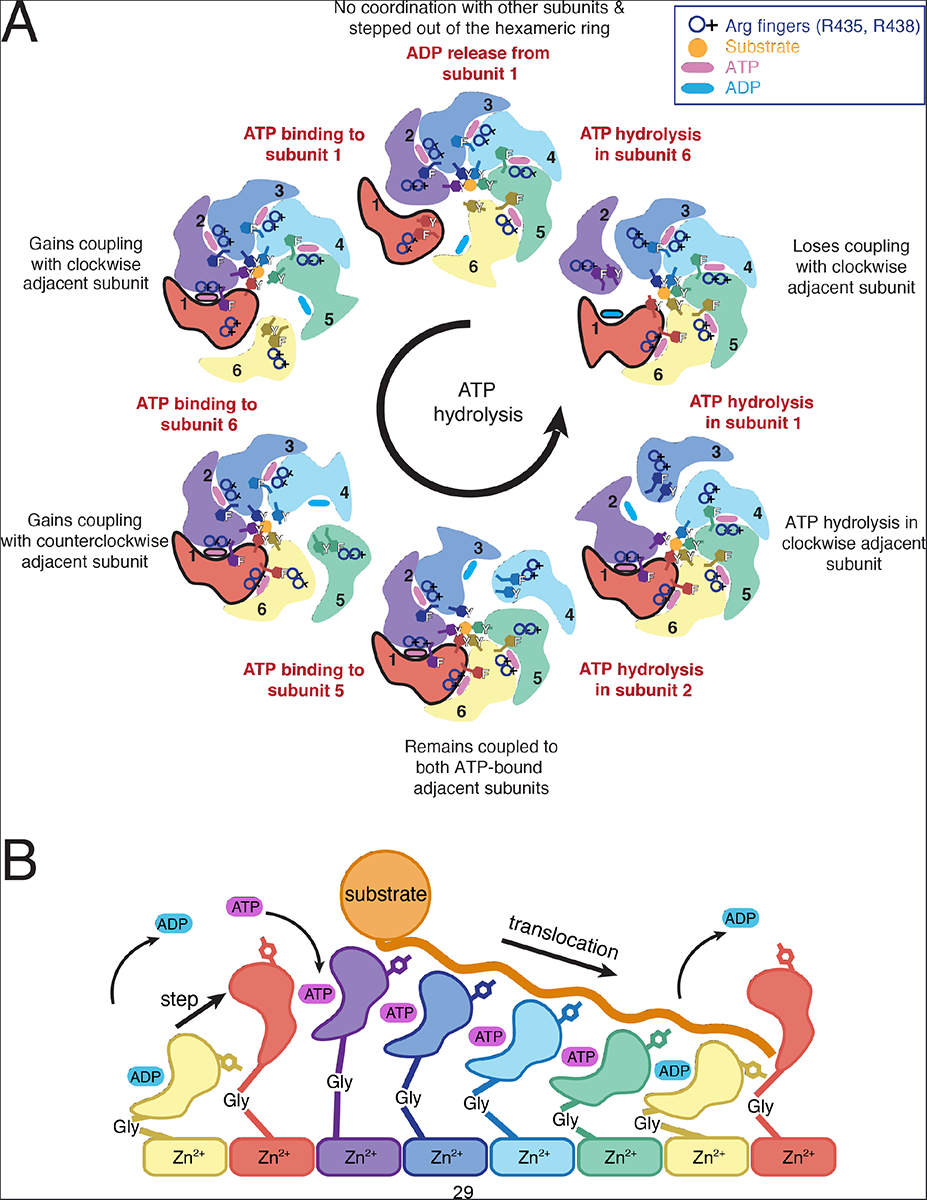
Model for ATP-driven substrate translocation into the proteolytic chamber of YME1. **A)** Cartoon representation of the ATP hydrolysis cycle that proceeds sequentially in a counterclockwise manner around the hexameric ATPase ring. We show how the coordinated nucleotide-dependent changes in the hexamer affect the subunit shown in red throughout the cycle. ATP and ADP are represented as blue and pink ovals at the interprotomer interface, and an orange circle depicts the substrate trapped in the central pore and coordinated by the pore-loops. **B)** Scheme of the progressive conformational rearrangements of a given subunit as it progresses through a single ATP-hydrolysis cycle (from left to right): ADP release, ATP-binding, ATP-bound states, ATP-hydrolysis, ADP-state, and ADP release. Each nucleotide state is highlighted by a different color. The yellow and red subunits are shown twice to highlight the complete cycle.

This tightly coordinated ATP-hydrolysis cycle and its allosteric relation to pore-loop conformation enables a constant grip on substrate as it is threaded through the central pore in a stepwise manner (Figure 5B). The ATP-bound subunits bind substrate, whereas the ADP-bound subunit releases the substrate at the lowest position of the spiral staircase, and the nucleotide-free subunit transitions to the highest position of the spiral staircase, where it reattaches to the substrate upon ATP-binding (movies 1 and 3). The unfolded substrate is translocated in this fashion toward the negatively charged proteolytic chamber, where it is positioned for cleavage at the zinc-coordinated active site of the immutable planar protease domains. Importantly, the large-scale motions of the ATPase closed spiral staircase observed in our structure are accommodated over a planar protease ring able to proceed without disrupting the protease architecture through a glycine residue within the linker connecting these domains.

### Concluding remarks

We determined the first high-resolution structure of the soluble domains of a mitochondrial IM AAA+ protease, providing a structural framework that explains decades of biochemical and genetic investigation of the activity and regulation of this class of unfoldase (Karata 2001, Yamada-Inagawa 2003). Furthermore, our asymmetric substrate-bound YME1 structure containing four ATP, one ADP-like, and one apo-like subunit is in agreement with previous biochemical data describing other unfoldases, including ClpX, PAN, and HslU, which showed a maximum of four ATP molecules and three coexisting functional subunit classes per hexamer (Hersch 2005, Horwitz 2007, Yakamavich 2008). Our model for ATP-driven substrate translocation incorporates and explains a number of previously identified mechanistic features from a wide variety of AAA+ ATPases, including direct interactions of the aromatic pore-loop residues with highly diverse substrates (Schlieker 2004, Siddiqui 2004, Hinnerwisch 2005, Park 2005, Nyquist and Martin 2014). In fact, we propose that the intercalation of pore loop tyrosines into the substrate polypeptide observed here is likely to be conserved across AAA+ unfoldases, as they enable substrate translocation independent of sequence. The presence of an additional tyrosine within the AAA+ staircase appears to be unique to YME1 (Supplementary Figure 6A), suggesting that increased hydrophobicity of the central pore may have evolved to improve the enzyme’s grip on greasy regions located within the numerous endogenous transmembrane substrates previously reported to be pulled out of the membrane by YME1 (Leonhard 2000, Tatsuta 2007, Augustin 2009). Furthermore, the increased structural flexibility of YME1 oligomers observed in the absence of nucleotide, which results from the loss of inter-subunit coordination, provides an explanation for the stress-induced proteolytic degradation of YME1L1 associated with reduced nucleotide levels that can impact recovery from ischemic reperfusion in mice (Baburamani 2015, Rainbolt 2015, Rainbolt 2016). The atomic description of how the AAA+ ATPase domains engage substrates will guide future studies aimed at unveiling the mechanisms that endow mitochondrial IM AAA+ proteases to act either as site-specific proteases or through processive degradation, and thereby regulate mitochondrial activity and morphology. These mechanisms are likely to be conserved given the high sequence similarity of these quality control complexes (Supplementary Figure 6A).

Recently solved substrate-bound structures of other AAA+ unfoldases have all revealed a staircase-like configuration of the ATPase subunits, leading to similar proposals for sequential nucleotide-driven translocation mechanisms (Gates 2017, Monroe 2017, Ripstein 2017). Although a certain degree of stochasticity cannot be discounted, the observed spiraling organization of the ATPase domains in these this and other substrate-bound structures of AAA+ unfoldases structures strongly supports a conserved, tightly coordinated ATP-hydrolysis cycle as the main driver of AAA+ ATPase activity. Furthermore, we confirm that three coexisting nucleotide states give rise to the ATPase staircase. We describe a new mechanism of allostery coordinating nucleotide state with substrate binding in unprecedented detail, and propose conservation across other type 1 ATPases, including the AAA+ ATPases of the 26S proteasome, which share over 35% sequence identity and show remarkable structural conservation of the closed spiral staircase configuration, as well as the key residues identified in YME1 (Supplementary Figure 6A,B). Indeed, the striking structural similarity between a yeast mitochondrial ATPase and a functionally related human cytosolic ATPase suggests that our proposed hydrolysis and translocation mechanism may be conserved throughout eukaryotic unfoldases and could help explain many aspects of the 26S proteasome translocation mechanism, which, despite many high resolution structures, remain unexplained (Budenholzer 2017).

## Materials and Methods

### Cloning and purification

A plasmid containing ^hex^YME1^WB^ was generated as previously described (Rampello and Glynn 2017). All additional variants of ^hex^YME1 were generated by site-directed mutagenesis using wild-type ^hex^YME1 as a template. All ^hex^YME1 variants were expressed and purified as previously described (Shi 2016, Rampello and Glynn 2017) with the following alterations. Proteins used in cryo-EM studies were buffer exchanged immediately prior to size exclusion chromatography into a buffer containing 20mM Tris-HCl (pH 8.0), 300mM NaCl, 2mM EDTA, 10% glycerol, and 1mM DTT. Proteins were then applied to a Superose 6 Increase column (GE Healthcare) pre-equilibrated in the same buffer. The peak corresponding to hexameric ^hex^YME1 was pooled, and buffer exchanged into a buffer containing 20mM Tris HCl (pH 8.0), 300mM NaCl, 15mM MgCl_2_, 10% glycerol, and 1mM DTT. Size exclusion was repeated using a Superose 6 increase column pre-equilibrated in a buffer containing 25mM HEPES (pH 8.0), 100mM KCl, 5% glycerol, 5mM MgCl_2_, and 1mM DTT. Fractions containing target proteins were pooled, concentrated and flash frozen prior to storage at -80 °C.

Sequence encoding I27 and I27^CD^ (Rampello and Glynn 2017) were subcloned into a modified pET His6 SUMO vector (2S-U) (15) (2S-U-I27; 2S-U-I27^CD^) and the T10 degron sequence was appended to the N-terminus of both constructs by PCR. All I27 variants were expressed in *E. coli* strain BL21-CodonPlus (Novagen) and purified as described for ^hex^YME1 substrates (Rampello and Glynn 2017).

### Biochemical assays

All ATPase and protein degradation assays were performed as previously described (Rampello and Glynn 2017). ATPase assays were carried out at 30 ^0^C and contained 0.2µM enzyme. Protein degradation reactions were carried out at 30^0^C and contained 20µM substrate, 0.5µM enzyme, and 5mM ATP regeneration system. Substrate degradation was visualized on 12% SDS-PAGE stained with Coomassie Blue R-250. Loss of full-length substrate band intensities were quantified using ImageJ (Schreiner 2012) and initial degradation rates calculated from at least five time points in the linear range. All intensities were normalized to 100% full-length substrate at 0 seconds (Gur 2012).

### Sample preparation for electron microscopy

For negative stain electron microscopy analysis of nucleotide induced conformational changes, 0.1 µM ^hex^YME1 was incubated for 5 minutes on ice in 25 mM HEPES pH 8, 100 mM KCl, 5 mM MgCl_2_, 1 mM DTT in the absence or presence of nucleotide (1 mM ATP or ADPALF_X_). After incubation, 4 µl of the sample were applied onto plasma cleaned 400 mesh Cu-Rh maxtaform grids (Electron Microscopy Sciences) coated with a thin layer of amorphous carbon. The grids were immediately stained with 2% (w/v) uranyl formate solution, and blotted to dryness.

After extensive cryo-EM screening to overcome strong preferred orientation, 0.76 mg/ml ^hex^YME1^WB^ was incubated on ice for 5 minutes in 25 mM HEPES pH 8. 100 mM KCl, 5 mM MgCl_2_, 1 mM DTT, 1 mM ATP and 0.05% Lauryl Maltose Neopentyl Glycol (LMNG, Anatrace). 3 µl of the sample were applied onto plasma cleaned UltrAuFoil Holey Gold Films R1.2/1.3, 300 mesh (Quantifoil) and immediately vitrified by plunge freezing in liquid ethane slurry at -180 °C. The entire procedure was performed using Vitrobot (Thermo Fisher) at 4 °C and 100% humidity.

### Electron microscopy data acquisition

Negative-stain EM micrographs were collected on a Tecnai Spirit (Thermo Fisher) transmission electron microscope operated at 120kV with a Lab6 filament using the Leginon automated acquisition software (Suloway 2005). Images were collected using an F416 CMOS 4Kx4K-pixel camera (TVIPS) at a nominal magnification of 52000x and a pixel size of 2.05 Å/pixel at specimen level. For each condition, 100,000 particles were picked from approximately 200 micrographs with an electron dose of 20 e^-^/Å^2^ using a defocus range of 0.5-1.5 µm.

Cryo-EM data were collected on a Thermo Fisher Titan Krios transmission electron microscope operating at 300keV and micrographs were acquired using a Gatan K2 Summit direct electron detector, operated in electron counting mode applying a total dose of 60 e^-^/Å^2^ as a 28-frame dose-fractionated movie during a 7s exposure. Leginon data collection software was used to collect 6098 micrographs at 29,000x nominal magnification (1.026 Å/pixel at the specimen level) and nominal defocus range of -1.2 to -2.5 µm.

### Image processing

For all negative stain data, the contrast transfer function was determined with CTFFind4 (Rohou and Grigorieff 2015) and particles were picked using Difference of Gaussians (DoG)-based automated particle picker (Voss 2009), both implemented in the Appion processing pipeline (Lander 2009). Final stacks of approximately 100000 particles were generated using RELION 1.4 (Scheres 2012) at a 144x144 box size, binned by a factor of 2 for processing. 2D classes were obtained using Reference free 2D alignment in Relion 2.0, but attempts to generate a reliable 3D reconstruction failed for all states, with the exception of the ATP-bound state.

During cryo-EM data collection, micrograph frames were aligned using MotionCorr2 (Zheng 2017), implemented in the Appion workflow (Lander 2009). CTF parameters were estimated with CTFFind4 (Rohou and Grigorieff 2015) and only micrographs with confidence values above 95% were further processed. Particles were picked with the FindEM template-based particle picker (Roseman 2004), using the negative stain 2D classes as templates. An initial 2,285,499 particle stack was created using a 256 pixel box size, which was scaled down by a factor of 4 using RELION 1.4 (Scheres 2012). 2D classification of these particles was performed in RELION 2.0 (Kimanius 2016), and only averages showing high resolution features were retained. The negative stain reconstruction was low pass filtered to 60 Å and used as an initial model for 3D refinement of 1,792,531 particles in RELION 2.0. The x & y shifts from this refinement were used to re-extract the centered, unbinned particles using a box size of 256 pixels. These particles refined to a reported resolution of 3.5 Å by FSC=0.143, but the resolution was severely anisotropic, and the reconstruction exhibited artifacts from preferred orientation. The particles from this reconstruction were then sorted by classification without alignment into five classes, two of which, accounting for 2% of particles each, displayed high-resolution features that did not display anisotropic resolution artifacts. The 62,917 particles from these classes were merged, refined, and post-processed to produce a final reconstruction with an estimated resolution of 3.4 Å by gold standard FSC at 0.143.

The cryo-EM density of the step subunit was poorly resolved in this reconstruction, so a soft edged 3D mask was generated to encompass the step subunit and used to “continue” the RELION refinement. Refinement of the masked step subunit improved the quality of the map in this region (Supplementary Figure 2D), with a reported resolution of 3.7 Å by FSC at 0.143.

### Atomic model building and refinement

A homology model was generated using the structure of a subunit of FtsH as a starting point. This initial model was split into the small and large domains of the ATPases and the protease and rigid body fit into the mrc density of each of the subunits. The structure of one of the subunits was refined using Phenix and COOT. Six copies of this atomic model were generated and each split into the 3 domains and rigid body fit into each subunit (step subunit density was used to build the corresponding atomic model). Further refinement in Phenix and COOT of the hexameric atomic model was performed using the nucleotide and coordinating metals as further restrictions. This refined model served as a starting point to generate 200 models in Rosetta and the top 5 scoring models were selected for further refinement in Phenix and COOT. The same procedure was followed for the step subunit atomic model using the step subunit density. Top 5 step subunit models were included in the pdb files of the top 5 models for the rest of the hexamer, respectively, resulting in the final 5 atomic models deposited. Poly-Alanine peptide was fit into the additional density using COOT. UCSF Chimera was used to generate the figures.

## Data and software availability

Electron microscopy maps, including sharpened, unsharpened, focused classification of the step subunit, and all associated half maps, are deposited to the Electron Microscopy Data Bank under accession number EMD-7023. Five atomic models that equally represent the EM density have been deposited as a multi-model entry at the Protein Data Bank under accession number PDB ID: 6AZ0.

## Acknowledgements

We thank Jean-Christophe Ducom at The Scripps Research Institute High Performance Computing for computational support, and Bill Anderson at The Scripps Research Institute electron microscopy facility for microscope support. We thank Bojian Ding for helpful discussions on sample preparation. C.P. is supported by an American Heart Association predoctoral fellowship. G.C.L. is supported as a Searle Scholar, a Pew Scholar, and by the National Institutes of Health (NIH) DP2EB020402. Computational analyses of EM data were performed using shared instrumentation funded by NIH S10OD021634 to G.C.L. A.J.R was supported by NIH training grant T32GM008468. A.J.R., C.J.G, and S.E.G. are supported by NIH R01GM115898. R.L.W. is supported by NIH R01NS095892.

## Author contributions

C.P. performed all cryo-EM structure determination, model building and refinement, mechanistic interpretation, and wrote the manuscript. C.P. and M.S. performed the negative stain EM experiments. A.J.R. and C.J.G. created all constructs and purified proteins. A.J.R. performed all biochemical experiments. All authors contributed to the experimental design and editing of the manuscript.

## REFERENCES

Augustin, S., F. Gerdes, S. Lee, F. T. Tsai, T. Langer and T. Tatsuta (2009). “An intersubunit signaling network coordinates ATP hydrolysis by m-AAA proteases.” Mol Cell 35(5): 574-585.

Baburamani, A. A., C. Hurling, H. Stolp, K. Sobotka, P. Gressens, H. Hagberg and C. Thornton (2015). “Mitochondrial Optic Atrophy (OPA) 1 Processing Is Altered in Response to Neonatal Hypoxic-Ischemic Brain Injury.” Int J Mol Sci 16(9): 22509-22526.

Baker, M. J., T. Tatsuta and T. Langer (2011). “Quality control of mitochondrial proteostasis.” Cold Spring Harb Perspect Biol 3(7).

Beckwith, R., E. Estrin, E. J. Worden and A. Martin (2013). “Reconstitution of the 26S proteasome reveals functional asymmetries in its AAA+ unfoldase.” Nat Struct Mol Biol 20(10): 1164-1172.

Bieniossek, C., B. Niederhauser and U. M. Baumann (2009). “The crystal structure of apo-FtsH reveals domain movements necessary for substrate unfolding and translocation.” Proc Natl Acad Sci U S A 106(51): 21579-21584.

Budenholzer, L., C. L. Cheng, Y. Li and M. Hochstrasser (2017). “Proteasome Structure and Assembly.” J Mol Biol.

Burton, R. E., T. A. Baker and R. T. Sauer (2005). “Nucleotide-dependent substrate recognition by the AAA+ HslUV protease.” Nat Struct Mol Biol 12(3): 245-251.

Chang, C. W., S. Lee and F. T. F. Tsai (2017). “Structural Elements Regulating AAA+ Protein Quality Control Machines.” Front Mol Biosci 4: 27.

DeLaBarre, B. and A. T. Brunger (2005). “Nucleotide dependent motion and mechanism of action of p97/VCP.” J Mol Biol 347(2): 437-452.

Enemark, E. J. and L. Joshua-Tor (2006). “Mechanism of DNA translocation in a replicative hexameric helicase.” Nature 442(7100): 270-275.

Gates, S. N., A. L. Yokom, J. Lin, M. E. Jackrel, A. N. Rizo, N. M. Kendsersky, C. E. Buell, E. A. Sweeny, K. L. Mack, E. Chuang, M. P. Torrente, M. Su, J. Shorter and D. R. Southworth (2017). “Ratchet-like polypeptide translocation mechanism of the AAA+ disaggregase Hsp104.” Science.

Gerdes, F., T. Tatsuta and T. Langer (2012). “Mitochondrial AAA proteases--towards a molecular understanding of membrane-bound proteolytic machines.” Biochim Biophys Acta 1823(1): 49-55.

Glynn, S. E., A. Martin, A. R. Nager, T. A. Baker and R. T. Sauer (2009). “Structures of asymmetric ClpX hexamers reveal nucleotide-dependent motions in a AAA+ protein-unfolding machine.” Cell 139(4): 744-756.

Graef, M. and T. Langer (2006). “Substrate specific consequences of central pore mutations in the i-AAA protease Yme1 on substrate engagement.” J Struct Biol 156(1): 101-108.

Graef, M., G. Seewald and T. Langer (2007). “Substrate recognition by AAA+ ATPases: distinct substrate binding modes in ATP-dependent protease Yme1 of the mitochondrial intermembrane space.” Mol Cell Biol 27(7): 2476-2485.

Gur, E., M. Vishkautzan and R. T. Sauer (2012). “Protein unfolding and degradation by the AAA+ Lon protease.” Protein Sci 21(2): 268-278.

Han, H., N. Monroe, J. Votteler, B. Shakya, W. I. Sundquist and C. P. Hill (2015). “Binding of Substrates to the Central Pore of the Vps4 ATPase Is Autoinhibited by the Microtubule Interacting and Trafficking (MIT) Domain and Activated by MIT Interacting Motifs (MIMs).” J Biol Chem 290(21): 13490-13499.

Hanson, P. I. and S. W. Whiteheart (2005). “AAA+ proteins: have engine, will work.” Nat Rev Mol Cell Biol 6(7): 519-529.

Hartmann, B., T. Wai, H. Hu, T. MacVicar, L. Musante, B. Fischer-Zirnsak, W. Stenzel, R. Graf, L. van den Heuvel, H. H. Ropers, T. F. Wienker, C. Hubner, T. Langer and A. M. Kaindl (2016). “Homozygous YME1L1 mutation causes mitochondriopathy with optic atrophy and mitochondrial network fragmentation.” Elife 5.

Hersch, G. L., R. E. Burton, D. N. Bolon, T. A. Baker and R. T. Sauer (2005). “Asymmetric interactions of ATP with the AAA+ ClpX6 unfoldase: allosteric control of a protein machine.” Cell 121(7): 1017-1027.

Hinnerwisch, J., W. A. Fenton, K. J. Furtak, G. W. Farr and A. L. Horwich (2005). “Loops in the central channel of ClpA chaperone mediate protein binding, unfolding, and translocation.” Cell 121(7): 1029-1041.

Horwitz, A. A., A. Navon, M. Groll, D. M. Smith, C. Reis and A. L. Goldberg (2007). “ATP-induced structural transitions in PAN, the proteasome-regulatory ATPase complex in Archaea.” J Biol Chem 282(31): 22921-22929.

Hoskins, J. R., S. M. Doyle and S. Wickner (2009). “Coupling ATP utilization to protein remodeling by ClpB, a hexameric AAA+ protein.” Proc Natl Acad Sci U S A 106(52): 22233-22238.

Karata, K., C. S. Verma, A. J. Wilkinson and T. Ogura (2001). “Probing the mechanism of ATP hydrolysis and substrate translocation in the AAA protease FtsH by modelling and mutagenesis.” Mol Microbiol 39(4): 890-903.

Kenniston, J. A., T. A. Baker, J. M. Fernandez and R. T. Sauer (2003). “Linkage between ATP consumption and mechanical unfolding during the protein processing reactions of an AAA+ degradation machine.” Cell 114(4): 511-520.

Kimanius, D., B. O. Forsberg, S. H. Scheres and E. Lindahl (2016). “Accelerated cryo-EM structure determination with parallelisation using GPUs in RELION-2.” Elife 5.

Koppen, M. and T. Langer (2007). “Protein degradation within mitochondria: versatile activities of AAA proteases and other peptidases.” Crit Rev Biochem Mol Biol 42(3): 221-242.

Lander, G. C., E. Estrin, M. E. Matyskiela, C. Bashore, E. Nogales and A. Martin (2012). “Complete subunit architecture of the proteasome regulatory particle.” Nature 482(7384): 186-191.

Lander, G. C., S. M. Stagg, N. R. Voss, A. Cheng, D. Fellmann, J. Pulokas, C. Yoshioka, C. Irving, A. Mulder, P. W. Lau, D. Lyumkis, C. S. Potter and B. Carragher (2009). “Appion: an integrated, database-driven pipeline to facilitate EM image processing.” J Struct Biol 166(1): 95-102.

Leonhard, K., B. Guiard, G. Pellecchia, A. Tzagoloff, W. Neupert and T. Langer (2000). “Membrane protein degradation by AAA proteases in mitochondria: extraction of substrates from either membrane surface.” Mol Cell 5(4): 629-638.

Leonhard, K., J. M. Herrmann, R. A. Stuart, G. Mannhaupt, W. Neupert and T. Langer (1996). “AAA proteases with catalytic sites on opposite membrane surfaces comprise a proteolytic system for the ATP-dependent degradation of inner membrane proteins in mitochondria.” EMBO J 15(16): 4218-4229.

Leonhard, K., A. Stiegler, W. Neupert and T. Langer (1999). “Chaperone-like activity of the AAA domain of the yeast Yme1 AAA protease.” Nature 398(6725): 348-351.

Martin, A., T. A. Baker and R. T. Sauer (2008). “Pore loops of the AAA+ ClpX machine grip substrates to drive translocation and unfolding.” Nat Struct Mol Biol 15(11): 1147-1151.

Matyskiela, M. E., G. C. Lander and A. Martin (2013). “Conformational switching of the 26S proteasome enables substrate degradation.” Nat Struct Mol Biol 20(7): 781-788.

Monroe, N., H. Han, P. S. Shen, W. I. Sundquist and C. P. Hill (2017). “Structural Basis of Protein Translocation by the Vps4-Vta1 AAA ATPase.” Elife 6.

Nolden, M., S. Ehses, M. Koppen, A. Bernacchia, E. I. Rugarli and T. Langer (2005). “The m-AAA protease defective in hereditary spastic paraplegia controls ribosome assembly in mitochondria.” Cell 123(2): 277-289.

Nyquist, K. and A. Martin (2014). “Marching to the beat of the ring: polypeptide translocation by AAA+ proteases.” Trends Biochem Sci 39(2): 53-60.

Okuno, T., K. Yamanaka and T. Ogura (2006). “Characterization of mutants of the Escherichia coli AAA protease, FtsH, carrying a mutation in the central pore region.” J Struct Biol 156(1): 109-114.

Park, E., Y. M. Rho, O. J. Koh, S. W. Ahn, I. S. Seong, J. J. Song, O. Bang, J. H. Seol, J. Wang, S. H. Eom and C. H. Chung (2005). “Role of the GYVG pore motif of HslU ATPase in protein unfolding and translocation for degradation by HslV peptidase.” J Biol Chem 280(24): 22892-22898.

Potting, C., T. Tatsuta, T. Konig, M. Haag, T. Wai, M. J. Aaltonen and T. Langer (2013). “TRIAP1/PRELI complexes prevent apoptosis by mediating intramitochondrial transport of phosphatidic acid.” Cell Metab 18(2): 287-295.

Potting, C., C. Wilmes, T. Engmann, C. Osman and T. Langer (2010). “Regulation of mitochondrial phospholipids by Ups1/PRELI-like proteins depends on proteolysis and Mdm35.” EMBO J 29(17): 2888-2898.

Quiros, P. M., T. Langer and C. Lopez-Otin (2015). “New roles for mitochondrial proteases in health, ageing and disease.” Nat Rev Mol Cell Biol 16(6): 345-359.

Rainbolt, T. K., N. Atanassova, J. C. Genereux and R. L. Wiseman (2013). “Stress-regulated translational attenuation adapts mitochondrial protein import through Tim17A degradation.” Cell Metab 18(6): 908-919.

Rainbolt, T. K., J. Lebeau, C. Puchades and R. L. Wiseman (2016). “Reciprocal Degradation of YME1L and OMA1 Adapts Mitochondrial Proteolytic Activity during Stress.” Cell Rep 14(9): 2041-2049.

Rainbolt, T. K., J. M. Saunders and R. L. Wiseman (2015). “YME1L degradation reduces mitochondrial proteolytic capacity during oxidative stress.” EMBO Rep 16(1): 97-106.

Rampello, A. J. and S. E. Glynn (2017). “Identification of a Degradation Signal Sequence within Substrates of the Mitochondrial i-AAA Protease.” J Mol Biol 429(6): 873-885.

Ripstein, Z. A., R. Huang, R. Augustyniak, L. E. Kay and J. L. Rubinstein (2017). “Structure of a AAA+ unfoldase in the process of unfolding substrate.” Elife 6.

Rohou, A. and N. Grigorieff (2015). “CTFFIND4: Fast and accurate defocus estimation from electron micrographs.” J Struct Biol 192(2): 216-221.

Roseman, A. M. (2004). “FindEM--a fast, efficient program for automatic selection of particles from electron micrographs.” J Struct Biol 145(1-2): 91-99.

Scheres, S. H. (2012). “RELION: implementation of a Bayesian approach to cryo-EM structure determination.” J Struct Biol 180(3): 519-530.

Schlieker, C., J. Weibezahn, H. Patzelt, P. Tessarz, C. Strub, K. Zeth, A. Erbse, J. Schneider-Mergener, J. W. Chin, P. G. Schultz, B. Bukau and A. Mogk (2004). “Substrate recognition by the AAA+ chaperone ClpB.” Nat Struct Mol Biol 11(7): 607-615.

Schreiner, B., H. Westerburg, I. Forne, A. Imhof, W. Neupert and D. Mokranjac (2012). “Role of the AAA protease Yme1 in folding of proteins in the intermembrane space of mitochondria.” Mol Biol Cell 23(22): 4335-4346.

Shah, Z. H., G. A. Hakkaart, B. Arku, L. de Jong, H. van der Spek, L. A. Grivell and H. T. Jacobs (2000). “The human homologue of the yeast mitochondrial AAA metalloprotease Yme1p complements a yeast yme1 disruptant.” FEBS Lett 478(3): 267-270.

Shi, H., A. J. Rampello and S. E. Glynn (2016). “Engineered AAA+ proteases reveal principles of proteolysis at the mitochondrial inner membrane.” Nat Commun 7: 13301.

Siddiqui, S. M., R. T. Sauer and T. A. Baker (2004). “Role of the processing pore of the ClpX AAA+ ATPase in the recognition and engagement of specific protein substrates.” Genes Dev 18(4): 369-374.

Stiburek, L., J. Cesnekova, O. Kostkova, D. Fornuskova, K. Vinsova, L. Wenchich, J. Houstek and J. Zeman (2012). “YME1L controls the accumulation of respiratory chain subunits and is required for apoptotic resistance, cristae morphogenesis, and cell proliferation.” Mol Biol Cell 23(6): 1010-1023.

Suloway, C., J. Pulokas, D. Fellmann, A. Cheng, F. Guerra, J. Quispe, S. Stagg, C. S. Potter and B. Carragher (2005). “Automated molecular microscopy: the new Leginon system.” J Struct Biol 151(1): 41-60.

Suno, R., H. Niwa, D. Tsuchiya, X. Zhang, M. Yoshida and K. Morikawa (2006). “Structure of the whole cytosolic region of ATP-dependent protease FtsH.” Mol Cell 22(5): 575-585.

Tatsuta, T., S. Augustin, M. Nolden, B. Friedrichs and T. Langer (2007). “m-AAA protease-driven membrane dislocation allows intramembrane cleavage by rhomboid in mitochondria.” EMBO J 26(2): 325-335.

Tatsuta, T. and T. Langer (2008). “Quality control of mitochondria: protection against neurodegeneration and ageing.” EMBO J 27(2): 306-314.

Voss, N. R., C. K. Yoshioka, M. Radermacher, C. S. Potter and B. Carragher (2009). “DoG Picker and TiltPicker: software tools to facilitate particle selection in single particle electron microscopy.” J Struct Biol 166(2): 205-213.

Vostrukhina, M., A. Popov, E. Brunstein, M. A. Lanz, R. Baumgartner, C. Bieniossek, M. Schacherl and U. Baumann (2015). “The structure of Aquifex aeolicus FtsH in the ADP-bound state reveals a C2-symmetric hexamer.” Acta Crystallogr D Biol Crystallogr 71(Pt 6): 1307-1318.

Wai, T., J. Garcia-Prieto, M. J. Baker, C. Merkwirth, P. Benit, P. Rustin, F. J. Ruperez, C. Barbas, B. Ibanez and T. Langer (2015). “Imbalanced OPA1 processing and mitochondrial fragmentation cause heart failure in mice.” Science 350(6265): aad0116.

Wang, K., M. Jin, X. Liu and D. J. Klionsky (2013). “Proteolytic processing of Atg32 by the mitochondrial i-AAA protease Yme1 regulates mitophagy.” Autophagy 9(11): 1828-1836.

Yakamavich, J. A., T. A. Baker and R. T. Sauer (2008). “Asymmetric nucleotide transactions of the HslUV protease.” J Mol Biol 380(5): 946-957.

Yamada-Inagawa, T., T. Okuno, K. Karata, K. Yamanaka and T. Ogura (2003). “Conserved pore residues in the AAA protease FtsH are important for proteolysis and its coupling to ATP hydrolysis.” J Biol Chem 278(50): 50182-50187.

Zhang, X. and D. B. Wigley (2008). “The ‘glutamate switch’ provides a link between ATPase activity and ligand binding in AAA+ proteins.” Nat Struct Mol Biol 15(11): 1223-1227.

Zheng, S. Q., E. Palovcak, J. P. Armache, K. A. Verba, Y. Cheng and D. A. Agard (2017). “MotionCor2: anisotropic correction of beam-induced motion for improved cryo-electron microscopy.” Nat Methods 14(4): 331-332.

